# Impaired glycolysis in aged endothelial cells is associated with reduced neovascularisation upon tissue ischemia

**DOI:** 10.1101/2024.01.26.577495

**Authors:** Kevin Kiesworo, Thomas Agius, Arnaud Lyon, Michael R Macarthur, Jing Zhang, Sébastien Déglise, James F. Markmann, Korkut Uygun, Heidi Yeh, Katrien De Bock, Sarah J. Mitchell, Alejandro Ocampo, Florent Allagnat, Alban Longchamp

**Author notes:** Contributed equally (Florent Allagnat, Alban Longchamp).

## Abstract

The vascular system experiences an age-associated decline in tissue perfusion and response to ischemic diseases. The factors driving this age-associated neovascularization decline remain unclear. While old endothelial cells (ECs) adopt a pro-angiogenic gene expression profile, we observed a stark reduction in the proliferative capacity of old ECs, while migratory capabilities remain intact. This is paralleled by a drastic decline in glycolytic capacity and ATP production, which likely act as limiting factors to neovascularization. These findings may provide new strategies to restore EC function in aging, thereby improving organ resilience and extending health- and lifespan.

## INTRODUCTION

Aging leads to a global loss of tissue structure and function, significantly increasing the risk of chronic illnesses. This includes the deterioration of the vascular system, resulting in an increased incidence of cardiovascular disease and broader organ impairment^1–6^. Conversely, the reversal of vascular decline alleviates age-related diseases and extends lifespan^7–10^.

The vascular network responds to varying tissue demands through neovascularization and vascular regression. Neovascularization is driven by the endothelial cells (ECs) lining the vessel lumen through three distinct processes: angiogenesis, arteriogenesis, and vasculogenesis^11,12^.

Various age-associated changes have been observed in the vasculature^4^. First and foremost, aging is associated with a phenomenon termed ‘microvascular rarefaction’, where capillary density diminishes with age across various organs^13,14^. This process leads to a reduction of tissue perfusion, which results in low-grade ischemia at resting state^15^. Importantly, rarefaction precedes the appearance of cellular hallmarks of aging (e.g senescence) in the tissue^14^. Another key observation is the decline of the neovascularisation potential with age, as previously shown in murine models of ischemic injury^16–18^. This age-associated decline in neovascularisation has been attributed to changes in EC migration and proliferation capacity, changes in EC metabolism^19^, and increased EC senescence and apoptosis^20–23^.

The aged tissue microenvironment contributes to altered vascular function. Aged organisms feature a systemic chronic, sterile, low-grade inflammation, known as inflammaging^24,25^, which compromises vascular function, impairs EC metabolism and survival^4,26,27^. Furthermore, aged tissues generate higher levels of ROS and oxidative damage^28–30^, which impairs EC migration, proliferation and angiogenesis^31,32^.

To dissect the mechanism underlying the age-associated decline in neovascularisation capacity, we undertook an unbiased gene expression analysis and metabolic profiling of young and old ECs. We discovered that while old ECs expressed elevated levels of pro-angiogenic genes at baseline, they may be ultimately restrained by dysfunctional metabolism, leading to reduced energy production that could prove crucial for cell proliferation, and consequently, neovascularisation.

## METHODS

### Animals

3-months (young) and 18-months (aged) old male C57BL/6J mice were used for young/old mice experiments as indicated in the text. All mice were housed at standard housing conditions (22°C, 12 h light/dark cycle), with ad libitum access to water and regular diet (SAFE®150 SP-25 vegetal diet, SAFE diets, Augy, France). All animal experimentations conformed to the National Research Council: Guide for the Care and Use of Laboratory Animals (National Research Council (U.S.)^33^. All animal care, surgery, and euthanasia procedures were approved by the CHUV and the Cantonal Veterinary Office (SCAV-EXPANIM, authorization number, 3504, 3768).

### Hindlimb ischemia (HLI) model

The HLI model was performed as previously described^34^. Briefly, mice were anaesthetized with isoflurane (2.5% under 2.5 L O_2_) and body temperature maintained on a circulating heated pad. Following a 1 cm groin incision, the neurovascular pedicle was visualized under a microscope (Z2 Zoom Stereo Microscope; LW Scientific, Lawrenceville, GA). The femoral nerve and vein were separated from the femoral artery. The femoral artery was ligated proximally, and above both “proximal caudal femoral” and “superficial caudal epigastric” arteries^35^ allowing electrocoagulation of the left common femoral artery while sparing the vein and nerve. Buprenorphine (0.1 mg/kg Temgesic, Reckitt Benckiser AG, Switzerland) was provided before surgery, as well as a post-operative analgesic every 12h for 36 hours. Mice were euthanized under anesthesia by cervical dislocation and exsanguination at time points indicated. Muscles were either frozen in OCT for histology, or flash frozen directly in liquid nitrogen for molecular analyses.

To measure cell proliferation *in vivo*, EdU (A10044, ThermoFisher Scientific) was diluted in NaCl at a concentration of 2 mg/ml and 500 μg was injected via i.p. injection 16h before sacrifice. Mice were sacrificed at day 5 post-HLI; ischemic muscles were placed in OCT and frozen in liquid nitrogen vapor.

### Laser Doppler perfusion imaging (LDPI)

Laser Doppler perfusion imaging (LDPI) was performed as described previously^34^. Briefly, mice were kept under isoflurane anesthesia, and body temperature maintained on a circulating heated pad. Once unconscious, we subjected the mouse hindlimbs to the LDPI (moorLDI2-HIR; Moor Instruments) system with a low-intensity (2 mW) laser light beam (wavelength 632.8 nm). Hindlimb blood flow was recorded as a 2D color-coded image, with a scan setting of 2 ms/pixel. Blood flow recovery was monitored at baseline, d0 (immediately post-surgery), d1, d3, d5, d7, d10, and d14. LDPI intensity of the ischemic foot was normalized to the corresponding contralateral foot and expressed as ratio between the ischemic/non-ischemic limb.

### Immunohistochemistry (IHC)

IHC was performed on 10 µm frozen sections of gastrocnemius muscle. After 5 min fixation in PFA 4% and rinsing in PBS, immunostaining was performed as previously described^34,36^. Muscle sections were permeabilized in PBS supplemented with 2 wt.% BSA and 0.1 vol.% Triton X-100 for 30 min, blocked in PBS supplemented with 2 wt.% BSA and 0.1 vol.% Tween 20 for another 30 min, and incubated overnight in the primary antibody diluted in the same buffer. Subsequently, the slides were washed three times for 5 min in PBS supplemented with 0.1 vol.% Tween 20, and incubated for 1 h at room temperature with a mix of appropriate fluorescent-labeled secondary antibodies. The antibodies used are described in **Supplementary Table 1**. The slides were then washed and mounted in DAPI-containing Vectashield fluorescent mounting medium. EdU immunostaining was performed according to the manufacturer’s instructions (Click-iT™ Plus EdU Cell Proliferation Kit for Imaging, Alexa Fluor™ 594 dye, ThermoFischer). Sections were then scanned with a Zeiss Axioscan microscope. VE-cad positive area of whole muscle was quantified blindly using Fiji software (ver. 1.53t; http://fiji.sc/Fiji). Quantifications were expressed as a percentage of VE-cad-positive area to the total surface area of the gastrocnemius muscle.

### Reverse transcription and quantitative polymerase chain reaction (RT-qPCR)

Pulverized frozen gastrocnemius muscles were homogenized in Tripure Isolation Reagent (Roche, Switzerland), and total RNA was extracted as published^34^. After RNA Reverse transcription (Prime Script RT reagent, Takara), cDNA levels were measured by qPCR Fast SYBR™ Green Master Mix (Ref: 4385618, Applied Biosystems, ThermoFisher Scientific AG, Switzerland) in a Quant Studio 5 Real-Time PCR System (Applied Biosystems, ThermoFischer Scientific AG, Switzerland). DNA oligo primers used in our analyses are described in **Supplementary Table 2**.

### Blood sampling

Peripheral blood was collected from the mice tail vein into EDTA tubes, pre-operatively and at 2 days post-HLI. Blood plasma are harvested and frozen at –80 °C. Leukocyte and erythrocyte fraction were then treated with 10ml of erythrocyte lysis buffer (0.15M NH_4_Cl, 5.7×10^-3^ KH_2_PO_4_, 1×10^-4^ Na_2_EDTA) for 5 minutes. Leukocytes were then stored in Gibco™ Recovery™ Cell Culture Freezing Medium (12648010, Thermo Fisher Scientific, Zurich, Switzerland).

### Luminex

Plasma at baseline and day 2 post-HLI were assayed by Luminex assay for Cytokine and chemokine 26-Plex Mouse ProcartaPlex Panel 1 (EPX260-26088-901, ThermoFisher Scientific AG, Switzerland) according to the manufacturer’s instructions.

### CyTOF

All samples were barcoded and processed simultaneously for antibody staining (markers selected: CD45, CD3, CD4, CD8, CD11b, CD11c, CD19, CD24, CD25, CD44, CD62L, CD64, CD103, CD117, F4/80, Ly6C, Ly6G/C, MHCII, NK1.1, SiglecF, TCRb, FoxP3, B220, CD69, CD127, Tbet, KLRG1; these prevalidated and pretitrated metal-conjugated antibodies were purchased from Fluidigm). The sample was mixed and incubated for 30 min at room temperature. After incubation, the sample was washed twice with Maxpar Cell Staining Buffer. Cells were then fixed by incubating the sample with 1 mL of 1.6% paraformaldehyde for 10 min. Subsequently, cells were washed twice with Maxpar Perm-S Buffer and centrifuged for 10 min at 1000g. Cells were then resuspended in 400 µL of Maxpar Perm-S Buffer and incubated for other 30 min with cytoplasmic/secreted antibody cocktail (1:100 dilution for each antibody, final volume 800 µL). At the end of the incubation, cells were washed twice with Maxpar Cell Staining Buffer and stained overnight with Cell-ID Intercalator-Ir solution at the final concentration of 125 × 10−9 m. Prior to data acquisition, the samples were washed twice with Maxpar Cell Staining Buffer, resuspended with 2 mL of Maxpar water and filtered using a

### 0.22 µm cell strainer cap to remove possible cell clusters or aggregates

Stained cells were analyzed on a CyTOF2/Helios mass cytometer (Fluidigm, San Francisco, USA) at an event rate of 400 to 500 cells per second. Data files for each barcoded sample were concatenated using an in-house script. The data were normalized using Normalizer v0.1 MCR. Files were debarcoded using the Matlab Debarcoder Tool.

### Muscle EC isolation, bulk RNA-seq, and analysis

Mice were euthanized and gastrocnemius muscles from both hindlimbs were immediately dissected and placed in an Petri dish on ice. Muscles were minced with a scalpel until a homogeneous paste-like mash was formed. EC isolation from muscle performed as described above. Cell suspension was centrifuged at 500 g for 5 min at 4°C, then the pellet was washed with ice cold HBSS (+20% FBS) followed by a centrifugation at 400 g for 5 min in 4°C. Next, the cell pellet was re-suspended in antibody medium (EGM2 CC-3162, Lonza, Basel, Switzerland) with anti-mouse CD31 PE antibody (1:400) (553373, BD Biosciences, Basel, Switzerland) and anti-mouse CD45 PerCP antibody (1:400) (557235 BD Biosciences, Basel, Switzerland) and placed on ice for 20 min in the dark. Before sorting, the cell suspension was washed in FACS buffer (1xPBS+1%BSA) and centrifuged at 400G for 5 min, 4°C, then the washed cell pellet was re-suspended in FACS buffer containing cell viability dye, SYTOX blue (1:1000) (S34857, Thermo Fischer Scientific, Zurich, Switzerland). Viable ECs (CD31+, CD45-, SYTOX blue-) were sorted by a FACS Aria III (BD Bioscience) sorter. ECs were directly sorted (70μm nozzle) into 700 μl RNA lysis buffer, and RNA extraction was performed using RNeasy Plus Micro Kit (74034 QIAGEN). RNA quality test was performed by Agilent High Sensitivity RNA ScreenTape System (G2964AA). Samples with RNA Integrity Number (RIN)≥8.0 were further analyzed.

Bulk RNA-seq libraries were obtained following the Smartseq II recommended protocol. Libraries were sequenced on the Novaseq 6000 (Illumina) instrument and sequenced data were processed using Kallisto to generate a count file matrix for each individual sample. Samples were pooled together on a single dataset for downstream analysis and genes with one count or less across all samples were filtered out.

### Analysis of Bulk RNASeq

The bioinformatics platform OmicsPlayground v2.8.10 (BigOmics Analytics; Lugano, Switzerland) was used for data pre-processing, statistical computation and visualization^37^. Data pre-processing included filtering of genes based on variance, the expression across the samples, and the number of missing values. Only protein coding genes on non-sexual chromosomes have been kept. A tSNE plot was generated to visualise the similarity in the gene expression profile of individual samples. For gene-level testing and identification of differentially expressed genes (DEG), statistical significance was assessed using two independent statistical methods: voom and limma-no-trend. The maximum q-value of the two methods was taken as aggregate q-value, which corresponds to taking the intersection of significant genes. A volcano plot was generated to visualise these genes.

Gene expression have been normalized using logCPM normalization in the edgeR R/Bioconductor package. All the 500 DEGs were analyzed for functional enrichment of GO terms and pathways using STRING v11.5 (Search Tool for the Retrieval of Interacting Genes), an online network cluster tool. To identify the most relevant clustering modules in the Protein– Protein Interactions network, we performed module/cluster analysis using the Markov Cluster Algorithm (MCL) with an infiltration of 1.8, in Cytoscape. The different clusters were then colored, and geneset enrichment was performed using 3 databases: GO Biological process (BP), KEGG pathway and Reactome pathway. Terms with a False Discovery Rate (FDR) <0.05 were collected. The clusters without relevant pathway enrichment or non-interacting genes were ignored. Chord diagram with top pathways for each have been performed using GOChord and clusterProfiler packages in R (version 4.1.2).

The genes in each cluster were subjected to Transcription factor (TF)-target gene regulatory network analysis to predict common transcription. We assessed transcription factor binding motifs, which are enriched in the genomic regions of a query gene set and thus allow for the prediction of transcription factors using the iRegulon plugin (version 1.3). To predict the transcriptional regulator and transcription factors using iRegulon analysis, we used genes from each cluster as input genes for motif and track search in the cis-regulatory control elements of genes. The criteria set for motif enrichment analysis were as follows: identity between orthologous genes ≥ 0.0, FDR on motif similarity ≤ 0.001, and TF motifs with normalized enrichment score (NES) > 4. The ranking option for Motif collection was set to 10 K (9713 PWMs) and a putative regulatory region of 20kb centered around TSS (7 species) was selected for the analysis.

### Aortic Ring Sprouting Assay

Aortic ring assay was previously described^38^. Briefly, mouse thoracic aortas were isolated and cut into 1 mm-wide rings and embedded in Matrigel® (Corning) and incubated in full EGM2 medium (Lonza). Media was replaced every two days. For each sample, the length of eight sprouts at days 6 and 9, originating from the aorta were quantified using the Fiji software (ImageJ 1.53t) rom brightfield images taken at 2X using a Nikon TI-2 live-cell imaging microscope.

### Cell culture

Pooled human umbilical vein endothelial cells (HUVECs; Lonza) were maintained in EGM™-2 (Endothelial Cell Growth Medium-2 BulletKit™; Lonza) at 37°C, 5% CO2 and 5% O_2_ as previously described^36^. Early passage HUVECs are defined to be under p10, whereas late passage HUVECs are above p15.

To understand the contribution of VEGF on proliferation and migration, we incubated HUVECs with the VEGF inhibitors KI8751 (Tocris Bioscience; 100 nM) and ZM323881 (Sigma-Aldrich; 2 µM) for the duration of the assay.

To harvest primary ECs, mice were euthanized with its lungs immediately dissected and placed in an Petri dish on ice. Lungs were minced with a scalpel until a homogeneous paste-like mash was formed. Subsequently, the lungs were enzymatically digested in a buffer containing 2 mg/ml Dispase II (D4693, Sigma-Aldrich, Steinheim, Germany), 2 mg/ml Collagenase IV (17104019, ThermoFisher Scientific, Zurich, Switzerland), 2 mM CaCl2 and 2% BSA in PBS at 37°C for 10 min, with gentle shaking every 3 min. The reaction was stopped by immediately adding an equal volume of ice cold HBSS containing 20% FBS and the suspension was passed through a 70-μm cell strainer (#431751, Corning, New York, USA) then 40-μm cell strainer (#431750, Corning, New York, USA) to remove tissue debris. Cell suspension was centrifuged at 500G for 5 min at 4°C, then the pellet was washed with ice cold HBSS (+20% FBS) followed by a centrifugation at 400G for 5 min in 4°C. Next, the cell pellet were re-suspended in 90μl HBSS (+20% FBS) and incubate with 10μl CD31 microbeads (130-097-418, Miltenyi Biotec, Bergisch GladbacH, Germany) in a FACS tube for 15 mins on ice. Wash cells, before resuspending in 1ml buffer. The tube will be transferred onto an EasySep™ magnet (18000, Stemcell Technologies, Vancouver, Canada). At this point, the purified ECs are resuspended in EGM™-2 for culturing.

### Cell Migration Assay

HUVEC were grown to confluence in a 12-well plate and a scratch wound was created using a sterile p200 pipette tip as previously described^34^. Repopulation of the wound in presence of Mitomycin C was recorded by phase-contrast microscopy over 16 hours in a Nikon Ti2-E live-cell imaging microscope. The denuded area was measured at t=0h and t=10h after the wound, and quantified using the ImageJ software (ver. 1.53t; http://fiji.sc/Fiji). Data were expressed as a ratio of the healed area over the initial wound area.

### Cell Transmigration Assay

The Boyden chamber assay was used to investigate the ability of ECs to transmigrate across a matrix barrier towards a chemotactic stimulus. This was performed using a 24-well tissue culture plates with Fluoroblok^TM^ inserts containing 8-μm pore-size polycarbonate membrane (Corning, New York, USA). HUVECs were trypsinized, and resuspended in EBM™-2 (Endothelial Cell Growth Basal Medium-2, Lonza). 500 μl EGM-2 BulletKit™ culture medium (with full supplements) was added into the bottom wells. ECs were subsequently loaded into the upper wells (10^5^ cells in 500 μl). After 8 h in cell culture, calcein-AM (5 µg/mL, ThermoFisher Scientific) was added to the well to stain the cells on the outer surface of the membrane of the transwell. After 30 min and two washes with PBS, fluorescence was measured using a fluorescent plate reader (Synergy Mx, BioTek; λex = 485 nm; λem = 530 nm).

### Cell Proliferation Assay

HUVEC were grown at 80% confluence (5×10^3^ cells per well) on glass coverslips in a 24-well plate and starved overnight in serum-free medium (EBM-2, Lonza). They were then incubated for 24 hr in EGM2 containing 10µM BrdU. Immunostaining was performed on cells washed and fixed for 5 min in -20°C methanol, air-dried, rinsed in PBS and permeabilized for 10 min in PBS supplemented with 2% BSA and 0.1% Triton X-100. BrdU positive nuclei were automatically detected in ImageJ software (ver. 1.53f51; http://fiji.sc/Fiji) and normalized to the total number of DAPI-positive nuclei^36^.

### Flow Cytometry (Cell Cycle Assessment)

HUVECs were grown at 70% confluence (5×10^4^ cells per well), before trypsinization and collection. Subsequently, the HUVECs were washed in ice-cold PBS before fixation by dropwise addition of ice-cold 70% ethanol while slowly vortexing the cell pellet. Cells were fixed for 1 h at 4 °C, washed 3 times in ice-cold PBS and resuspended in PBS supplemented with 20 µg/mL RNAse A and 10 µg/mL DAPI. Flow cytometry was performed in a Cytoflex-S apparatus (Beckman Coulter GmbH, Krefeld, Germany).

### Seahorse

Glycolysis and Mitochondrial stress tests were performed on confluent HUVECs according to the manufacturer’s kits and protocols (Agilent Seahorse XF glycolysis stress test kit, Agilent Technologies, Inc.). 1 μM Oligomycin was used. Data were analyzed using the Seahorse Wave Desktop Software (Agilent Technologies, Inc., Seahorse Bioscience).

### Senescence-Associated Beta Galactosidase Staining

HUVECs were cultured on glass cover slips at 80% confluence. Cells were washed with PBS before fixing with neutral buffered 4% PFA for 3 mins at room temperature. After another wash, fixed cells were incubated for 12-16h at 37°C with staining solution for beta galactosidase (1mg/ml X-gal, 1X citric acid/sodium phosphate buffer pH 6.0, 5mM potassium ferricyanide, 5mM potassium ferrocyanide, 150mM NaCl, 2mM MgCl_2_). Subsequently, the coverslips were then washed once before mounting with 50% glycerol on microscope slides. Images were taken on Nikon TS100 light microscope.

### HUVEC incubation, metabolite extraction and measurements

EGM-2 media was replaced with DMEM media containing D-Glucose (U-^13^C_6_, CLM-1396-PK, Cambridge Isotope Laboratories Inc.) 3 hours prior to collection. Prior to harvest, labelled media was removed, and HUVECs were washed with PBS. Metabolite extraction from HUVECs was performed as rapidly as possible to minimize perturbations in metabolism, and the extracts were analyzed by LC–MS. Cell metabolism was quenched, and metabolites were extracted by quickly aspirating media from culture dishes and adding cold (–20 °C) 80:20 methanol/water. Then, cells were scraped off the culture dish surface. Note that, in all cases, quenching was performed without any washing steps that can perturb the metabolism.

Once obtained, all extracts were moved to Eppendorf tubes for centrifugation at 4 °C. The supernatants were dried and reconstituted in HPLC-grade water. These samples were analyzed by reversed-phase ion-pairing liquid chromatography coupled to a high-resolution orbitrap mass spectrometer with electrospray ionization in negative-ion mode and a resolution of at least 100,000 at m/z 200 (Exactive and Q Exactive, Thermo). The LC–MS data were analyzed using the Metabolomic Analysis and Visualization Engine (MAVEN) with analyte peaks identified by authenticated standards. Measured labeling fractions were corrected for the natural abundance of 13C, as well as the impurities in the labeled substrates.

### Statistical analyses

Data are displayed as means ± standard error of the mean (S.E.M.). Statistical significance was assessed in GraphPad Prism 9.1.0 using Student’s t-test, one-way or two-way ANOVA unless otherwise specified. A p-value of 0.05 or less was deemed statistically significant.

## RESULTS

### Reduced EC proliferation capacity impairs post-HLI neovascularisation response in aged mice

When compared to young (3 m.o.) mice, aged (18 m.o.) mice displayed reduced capillary density of the gastrocnemius, soleus and adductor muscles (Fig. 1A<u>, S1A</u>). To explore the effects of aging on vascular functionality, we utilised the murine hindlimb ischemia (HLI) model, which involves the unilateral ligation of the femoral artery, resulting in leg ischemia. When subjected to HLI, we observed an equivalent reduction in tissue perfusion in the ischemic hindlimb in both young and aged mice. However, the recovery of tissue perfusion was slower in the aged mice (Fig. 1B).

**Fig. 1:**
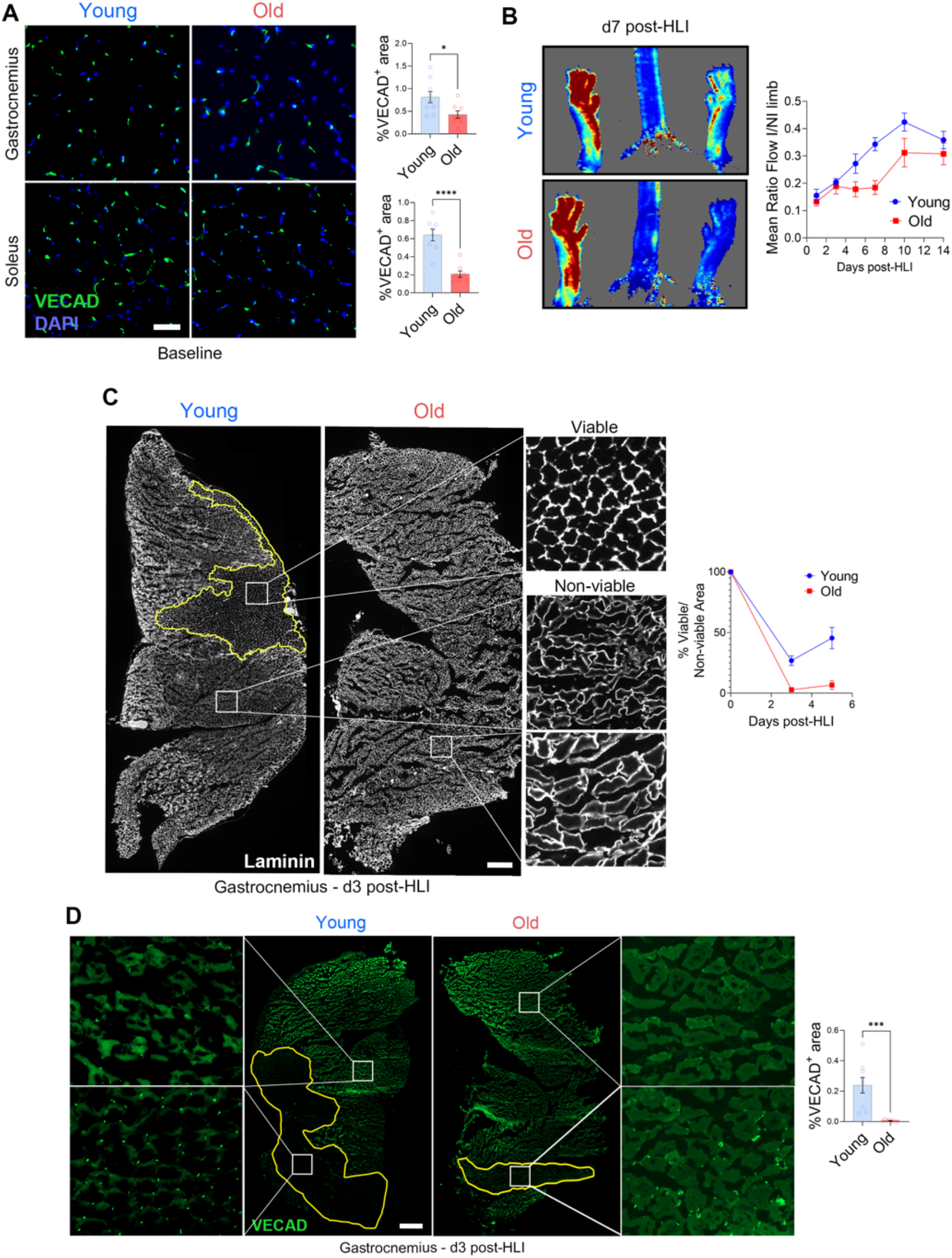
Aged mice undergo slower neovascularisation and suffer more muscle damage post-hindlimb ischemia. **(A)** Gastrocnemius and soleus muscle microvasculature at baseline as depicted by representative images (left) and quantification (right) of transverse sections stained with VE-cad. n=8-9 per group. Scale bar represents 50µm. **(B)** Superficial perfusion of mouse hindlimb measured by Laser Doppler Perfusion Imaging (LDPI). Results illustrated by the ratio of LDPI perfusion quantification in ischemic and non-ischemic limbs (left) and representative LDPI images (right). n=8 per group. **(C)** Representative transverse sections (left) and quantification (bottom right panel) of laminin staining of gastrocnemius muscle at 3 days post-HLI. n=4-8 per group. Scale bar represents 50µm. **(D)** Ischemic gastrocnemius muscle microvasculature at 3 days post-HLI as depicted by representative images (left) and quantification (right) of transverse sections stained with VE-cad. Areas within bordered white line represent areas with well-ordered, viable vasculature. n=8-9 per group. Scale bar represents 50µm. Data are expressed as mean ± S.E.M. ** p≤0.01, *** p≤0.001 and **** p≤0.0001 by Student’s t-test.

At day 3 and 5 post-HLI, we observed a larger area of disordered laminin patterning within the ischemic gastrocnemius muscle of aged mice, indicative of elevated tissue damage (d3: 9.3-fold, d5: 6.7-fold larger; Fig. 1C). Concurrently, a 40-fold reduction in VE-Cadherin (VE-Cad; EC marker) positive signal was observed at day 3 post-HLI on the ischemic gastrocnemius (Fig. 1D) and soleus muscles (<u>Fig. S1B</u>) of the aged mice. Within the damaged regions of the gastrocnemius and soleus muscles, we observed VE-Cad positive signal in the young mice, but not in the old mice (<u>Fig. S1C</u>), which may be indicative of active neovascularisation only in the young mice. Capillary density of aged mice gastrocnemius muscle remained lower at day 21 post-HLI (<u>Fig. S1D</u>).

Arteriogenesis, the development of collateral vessels from pre-existing blood vessels, contributes to vascular repair in response to HLI^39,40^. We did not observe differences in smooth muscle alpha-actin (α-SMA; arterial marker) positive signal and arteriole number in the adductor muscle between young and aged mice (<u>Fig. S1E</u>). However, we observed a 2.6-fold lower α-SMA-positive signal in the ischemic adductor muscle of the aged mice at day 5 post-HLI (Fig. 2A), suggesting decreased arteriogenesis.

**Fig. 2:**
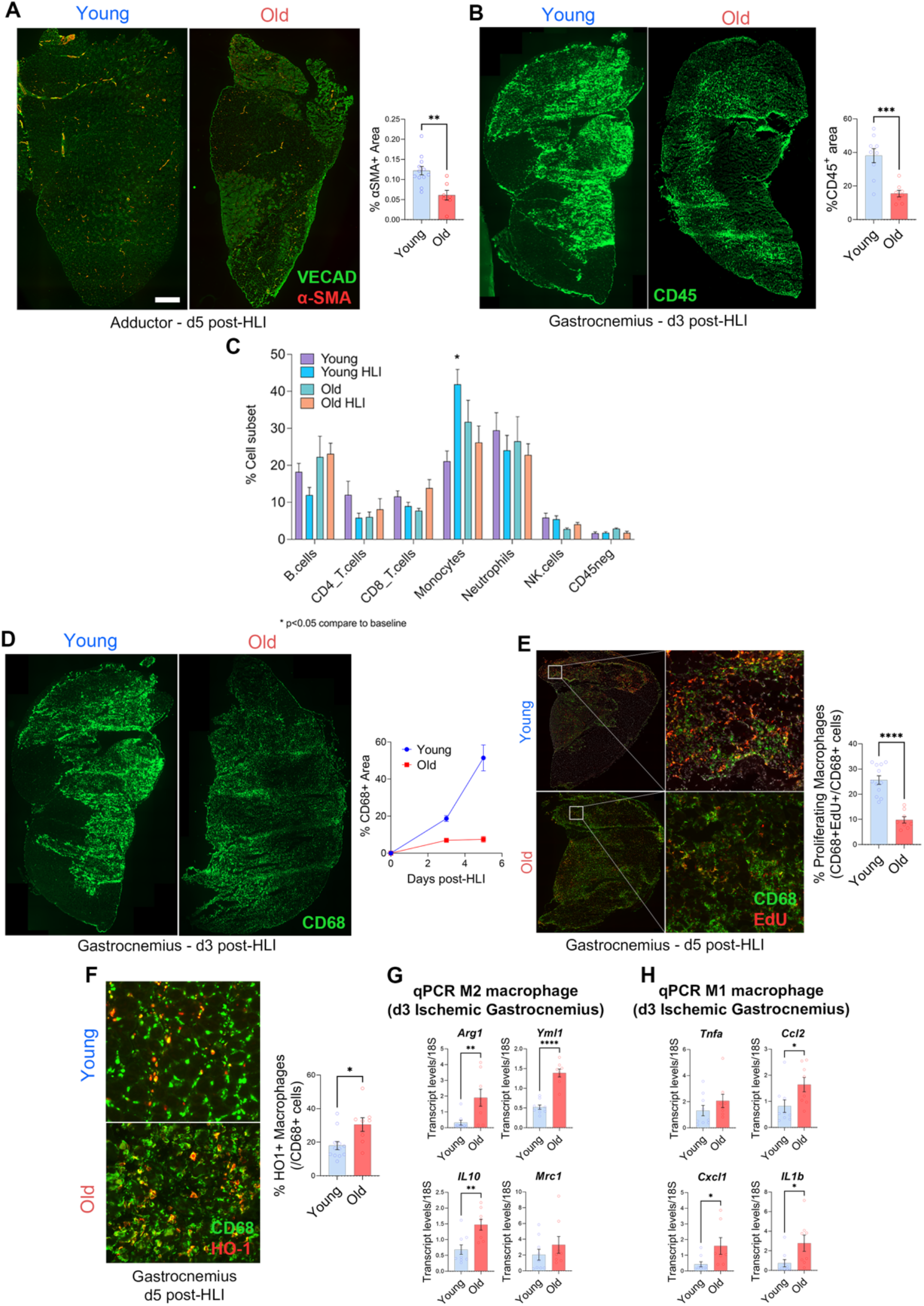
Arteriogenesis defect correlates with impairment in immune recruitment in aged mice muscle post-hindlimb ischemia. **(A)** Arterioles in the ischemic adductor muscle at 5 days post-hindlimb ischemia as depicted by representative images (left) and quantification (right) of transverse sections stained with VE-cad and α-SMA. n=8-12 per group. Scale bar represents 50µm. **(B)** Immune cell infiltration in gastrocnemius muscle at 3 days post-hindlimb ischemia as depicted by representative images (left) and quantification (right) of transverse sections stained with CD45. n=8-9 per group. Scale bar represents 150µm. **(C)** Frequencies of B cells, CD4+ T cells, CD8+ T cells, monocytes, neutrophils, NK and CD45-cells at baseline and at 2 days post-HLI. Cell frequency expressed as a percentage of total hematopoetic cells. **(D)** Macrophage infiltration in gastrocnemius muscle as depicted by representative images at 3 days post-hindlimb ischemia (left) and quantification at 3 and 5 days post-hindlimb ischemia (right) of transverse sections stained with CD68. n=8-10 per group. Scale bar represents 150µm. **(E)** Proliferating macrophages in gastrocnemius muscle at 5 days post-hindlimb ischemia as depicted by representative images (left) and quantification (right) of transverse sections stained with CD68 and EdU. n=8-10 per group. Scale bar represents 150µm. **(F)** M2-like macrophages in gastrocnemius muscle at 5 days post-hindlimb ischemia as depicted by representative images (left) and quantification (right) of transverse sections stained with CD68 and EdU. n=8-10 per group. Scale bar represents 150µm. **(G-H)** Quantitative real-time PCR gene expression analysis of (G) M2 macrophage (H) M1 macrophage markers in ischemic gastrocnemius muscle at 3 days post-hindlimb ischemia. All genes were normalised to 18S expression. n=8-9 per group. Data are expressed as mean ± S.E.M. * p≤0.05, ** p≤0.01, *** p≤0.001 and **** p≤0.0001 by Student’s t-test.

Immune cells play an important role in supporting arteriogenesis^41–43^, with monocytes homing towards the collateral arteries to promote remodelling through the paracrine effects of chemokines, metalloproteinases, and growth factors^44–46^. At day 3 post-HLI, infiltration of CD45^+^ haematopoetic cells (Fig. 2B) was reduced 2.5-fold in the ischemic gastrocnemius muscle of aged mice. To examine the age-related differences in the immune response to HLI, we analysed peripheral blood cells with mass cytometry (CyTOF), and performed an unsupervised clustering algorithm to distil multidimensional single-cell data (<u>Fig. S2</u>). At baseline, we observed no age-related differences in any of the annotated cell subsets. At day 2 post-HLI, the percentage of circulating monocytes was increased in young mice, but not in old mice. No significant differences were observed in other cell subsets post-HLI (Fig. 2C). CD68^+^ macrophages infiltration in the muscle was reduced 2.7 fold at day 3 and 6.9-fold at day 5 post-HLI in aged mice (Fig. 2D) Moreover, macrophage proliferation was reduced 2.6-fold in aged mice at day 5 post-HLI (Fig. 2E). During vascular repair following HLI, pro-inflammatory M1-like macrophages shift their phenotype to a pro-regeneration M2-like phenotype to support neovascularisation^47,48^. CD68^+^ HO1^+^ M2-like macrophages were increased 1.7-fold in the ischemic calf muscle of aged mice at day 5 post-HLI (Fig. 2F), suggesting an earlier shift toward pro-repair M2 macrophage in the old mice. However, qPCR analyses revealed significant increases in the expression of both M2-like macrophages markers (Arg1, Yml1, and IL10; Fig. 2G) and M1-like pro-inflammatory macrophages markers (Ccl2, IL1b, Cxcl1; Fig. 2H). Overall, age was associated with impaired macrophage infiltration but a larger proportion of macrophages shifted toward a pro-repair phenotype. This does not correlate with defective arteriogenesis and unlikely to contribute much to impaired neovascularisation. Thus, we then focused on EC proliferation and migration, and sprouting angiogenesis.

We observed a 6.4-fold reduction in the proportion of proliferating endothelial cells (EdU^+^Erg^+^) in the calf muscles of aged mice at day 5 post-HLI (Fig. 3A). Moreover, sprouting angiogenesis from aortic explants was reduced 2-fold in aged compared to young mice (Fig. 3B). To look further into this phenomenon, we utilised an *in vitro* model of EC aging comparing early (1-10) and late passages (>15) of primary human umbilical vein endothelial cells (HUVECs) (Fig. 3C). The late passage HUVECs showed a 8.6-fold larger proportion of β-Galactosidase-positive cells (Fig. 3D), a validated marker of cellular senescence^49^. While migratory capacity was maintained (Fig. 3E<u>, S3A</u>), late passage HUVECs displayed a 3.3-fold reduction in proliferation (Fig. 3F). Moreover, late passage HUVEC accumulated in the G1 phase and displayed increased level of the G1 phase marker Cyclin D1, whereas the S phase markers Cyclin E1 and Cyclin-dependent kinase 2 (CDK2) were reduced (Fig. 3G<u>; Fig. S3B</u>).

**Fig. 3:**
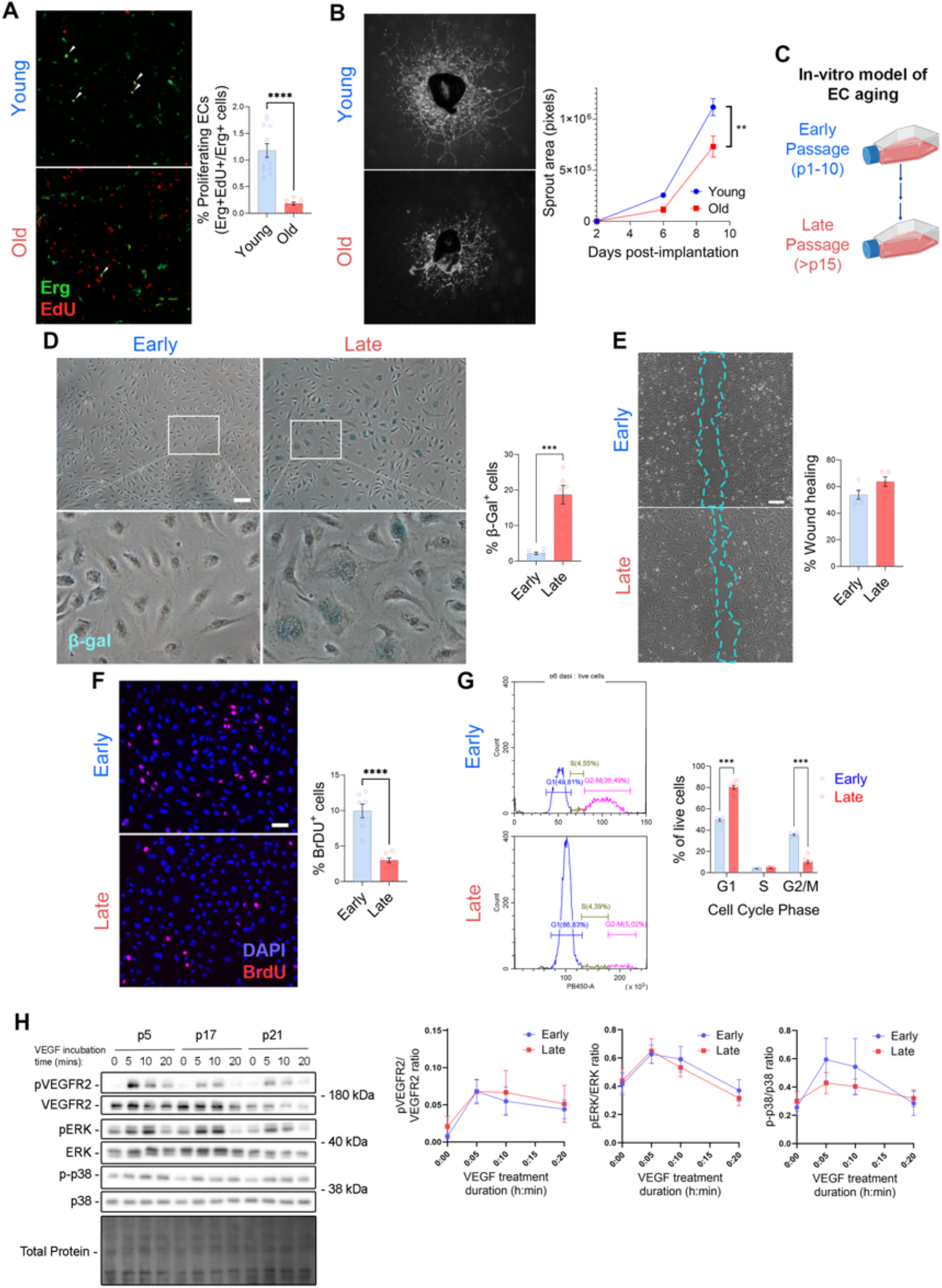
Impairment of EC proliferation results in neovascularisation defect in old mice. **(A)** Proliferating ECs in gastrocnemius muscle at 5 days post-hindlimb ischemia as depicted by representative images (left) and quantification (right) of transverse sections stained with Erg and EdU. n=8-10 per group. Scale bar represents 150µm. **(B)** Average area of microvessel sprouting from aortic ring explants from young or old mice incubated in full EGM2 media as represented by images at day 9 post-implantation in matrigel (left) and quantification (right). n=8-10 per group. Scale bar represents 100µm. **(C)** *In vitro* model of EC aging. HUVECs are continually passaged in culture; and early passage cells are defined as passages 1-10 and late passage cells are defined as beyond passage 15. **(D)** Senescent HUVECs as depicted by representative images (left) and quantification (right) of an early and late passage HUVEC monolayer stained with β-galactosidase. n=6 per group. Scale bar represents 150µm. **(E)** EC migration across scratch with representative image (left, 10x magnification) and quantification (right) of early and late passage HUVECs. n=5 per group. Scale bar represents 150µm. **(F)** Proliferating HUVECs as depicted by representative images (left) and quantification (right) of an early and late passage HUVEC monolayer stained with BrdU and DAPI. n=7 per group. Scale bar represents 150µm. **(G)** Cell cycle dynamics of early and late passage HUVECs, as described by a representative flow cytometry plot of DAPI expression against cell count (left) and quantification of the proportion of live cells within G1, S and G2/M phases of the cell cycle (right). n=4-7 per group. **(H)** Western blot on lysates of early and late passage HUVECs treated with 20nM VEGF for the duration of 0, 5, 10, and 20 minutes. P-VEGFR2, P-ERK and P-p38 levels normalized with total VEGF, ERK and p38 levels respectively. All proteins normalized to total protein stain. Data are expressed as mean ± S.E.M. * p≤0.05, ** p≤0.01, *** p≤0.001 and **** p≤0.0001 by Student’s t-test otherwise stated.

However, the protein expression of the cyclin-dependent kinase inhibitor p21 (also known as p21^WAF1/Cip1^), which promotes cell cycle arrest in response to a variety of stimuli, was also severely reduced, suggesting no direct induction of cell cycle arrest (<u>Fig. S3B</u>). There was no difference between early and late HUVEC in VEGF-induced phosphorylation of VEGFR2, ERK and p38 (Fig. 3H), indicating no impaired VEGF response in late passage HUVECs. In line with this observation, inhibition of VEGF signalling using two VEGFR inhibitors KI8751 and ZM323881 impacted cell migration, but not proliferation in early passage HUVECs (<u>Fig. S3C,D</u>). Thus, the defective cell proliferation in old ECs was not due to impaired VEGF response or cell cycle arrest.

### Aged ECs display altered gene expression in angiogenic and metabolic pathways

To understand the age-related impaired EC proliferation and vascular repair potential, we undertook an unbiased comparison of the gene expression profiles of ECs from the gastrocnemius muscle harvested from young and old mice by bulk RNASeq. Unsupervised clustering demonstrated a clear divergence in gene expression phenotype between young and old ECs (Fig. 4A) – with 500 differentially expressed genes (DEGs; 406 upregulated, 94 downregulated; Fig. 4B). Gene network analysis of DEGs revealed 12 clusters, 10 upregulated and 2 downregulated (Fig. 4C).

**Fig. 4:**
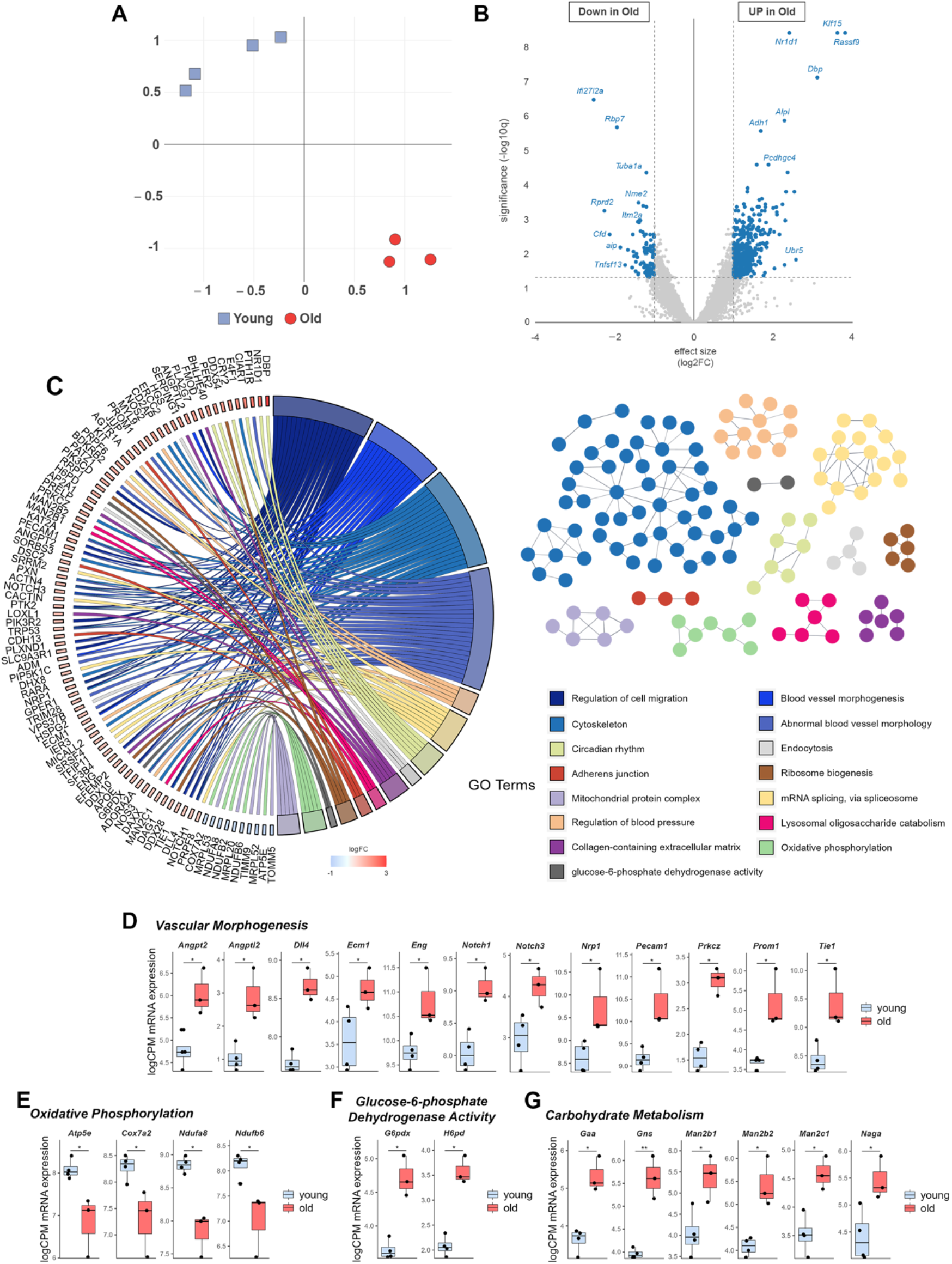
Old ECs exhibit an altered gene expression profile. **(A)** tSNE plot visualising the degree of similarity in gene expression profiles of individual samples of young (blue) and old (red) ECs. n=3-4 per group. **(B)** Volcano plot representing the identification of 500 genes that are differentially expressed with age in skeletal muscle ECs. n=3-4 per group. **(C)** STRING protein interaction networks functional enrichment analysis of differentially expressed genes (DEGs; left). Chord diagram representing the connections between DEGs and functional pathways, differentiated by colour (right). n=3-4 per group. **(D-G)** mRNA expression of genes associated with vascular morphogenesis (D), oxidative phosphorylation (E), glucose-6-phosphate dehydrogenase activity (F), carbohydrate metabolism (G), expressed as logarithm of counts per million reads (logCPM).

Interestingly, we observed an upregulation of a large gene cluster associated with pro-angiogenic vascular morphogenesis (Fig. 4D) in aged ECs. This cluster contains Notch1 and Dll4, genes central in tip-stalk cell speciation during angiogenesis^50–52^, Nrp1, a VEGF receptor vital for EC migration towards a VEGF gradient^53^, and the growth factor Angpt2, which enhances tip cell migration and vascular sprouting^54^, and Tie1, which is required for full activation of Tie2 by Angpt1/Angpt2^55,56^.

Aged ECs also exhibited alterations in metabolic gene expression levels. Genes involved in oxidative phosphorylation were downregulated (Fig. 4E), while genes associated with the pentose phosphate pathway (G6pdx and H6pd) and carbohydrate metabolism (e.g Gaa, Gns) were upregulated (Fig. 4F, G). These metabolic adaptations are reminiscent of EC metabolic reprogramming in the context of angiogenesis^57–63^.

### Age-associated glycolysis impairment in ECs may contribute to sub-optimal neovascularisation capacity

To further assess EC metabolism, we investigated the glycolytic capacity of EC by Seahorse. We found that both ECs harvested from aged mice and late passage HUVEC had severely decreased glycolytic capacity (Fig. 5A, B). To investigate the impact of passage number on HUVEC metabolic flux, HUVECs cultured for 3 hours in presence of uniformly labelled ^13^C_6_ were analysed by mass spectrometry. PCA analysis of early and late passage HUVEC revealed clearly distinct populations (Fig. 5C). Late passage HUVECs exhibited a reduction in ^13^C_6_ enrichment of glycolytic metabolites (Fig. 5D; hexose phosphate, fructose 1,6-biphosphate, 2-phosphoglyceric acid and lactate). Although ^13^C_6_ incorporation is reduced in hexose phosphate, we observe no passage differences in enrichment into the pentose phosphate pathway intermediate ribose-5-phosphate (Fig. 5E), an important precursor for nucleotide biosynthesis. Similarly, we do not observe passage differences in ^13^C_6_ incorporation into the serine biosynthesis pathway (Fig. 5F; 3-phosphohydroxypyruvate, phosphoserine, and serine), also important for nucleotide biosynthesis. While we observe a smaller percentage of ^13^C_6_ labelled acetyl-CoA and aconitate in late passage HUVECs, we observe an increase in ^13^C_6_ enrichment in the later intermediates of the TCA cycle (succinic, fumaric and malic acids) in late passage HUVECs (Fig. 5G). Lastly, we observed a reduction of ^13^C_6_ incorporation into ATP (Fig. 5H), signifying a reduced level of glucose-derived energy generation.

**Fig. 5:**
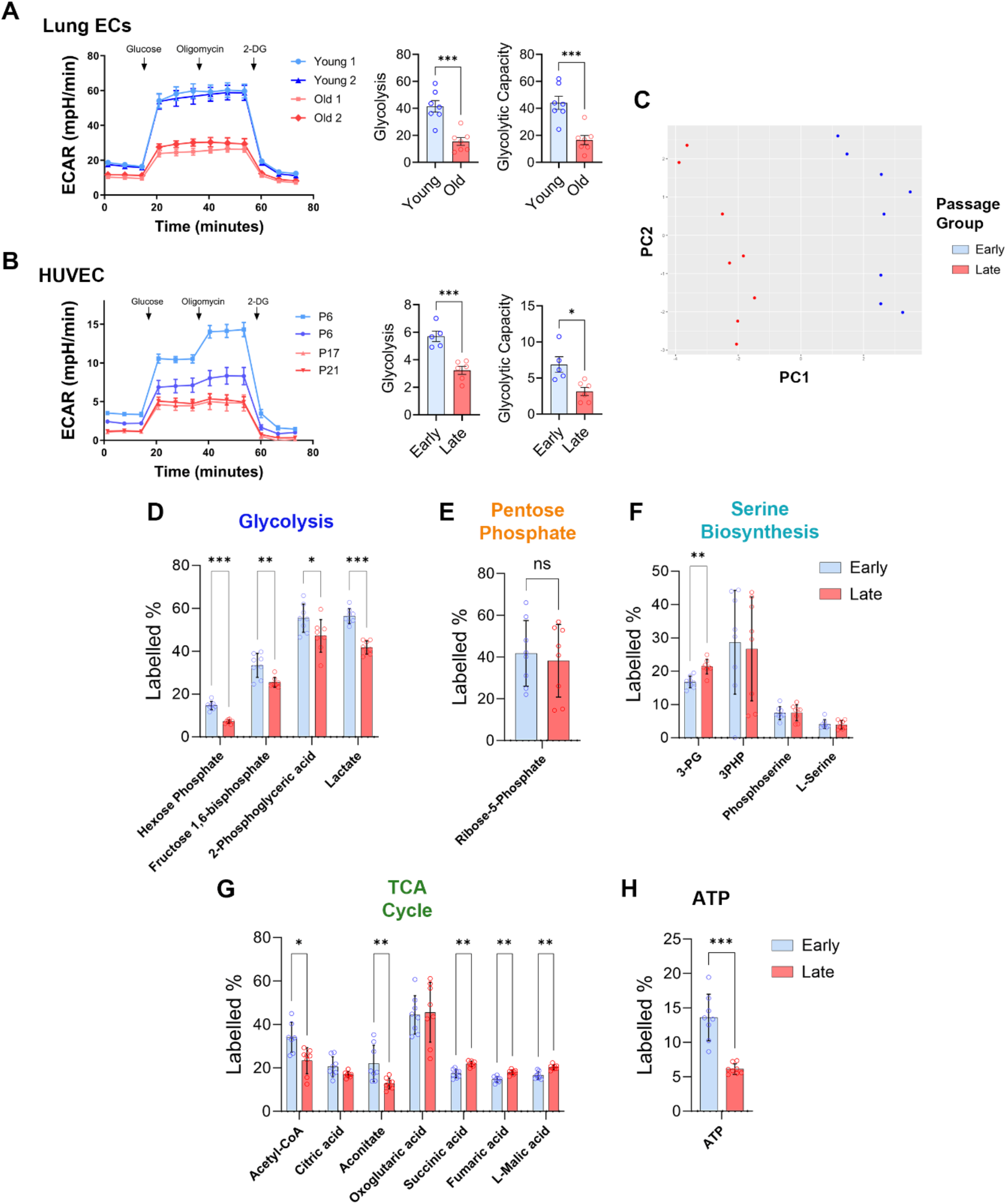
Glucose metabolism is impaired in old ECs and late passage HUVECs. **(A-B)** Glycolysis stress test in (A) ECs harvested from 3-or 18-month old mice, and (B) early and late passage HUVECs. Glycolysis is measured by extracellular acidification rate (ECAR). Data are expressed as mean ± S.E.M. * p≤0.05, ** p≤0.01, *** p≤0.001 and **** p≤0.0001 by Student’s t-test. **(C)** PCA clustering of total enrichment data from U-^13^C_6_-labelled glucose tracing experiment to investigate passage differences in baseline HUVEC intracellular metabolism. **(D-H)** ^13^C_6_ enrichment of metabolites within the (D) glycolysis, (E) pentose phosphate, (F) serine biosynthesis, and (G) TCA cycle pathways, and (H) ^13^C_6_ enrichment into ATP. Statistical significance determined through the Mann-Whitney test; * p < 0.05; ** p < 0.01; *** p < 0.001. n=8 per group.

## DISCUSSION

In this study, we investigated why old endothelial cells (ECs) exhibit diminished neovascularization capacity, a critical factor in understanding age-related vascular decline. Our research confirms the age-related decrease in tissue capillary density^14^ and impaired neovascularization response to ischemia^16,17^. We demonstrate that this impairment is likely linked to failure to proliferate secondary to metabolic dysfunction and ATP depletion in ECs, despite a gene expression profile characteristic of active proliferation and angiogenesis.

Neovascularization following hindlimb ischemia involves both arteriogenesis and angiogenesis. Arteriogenesis necessitates pro-resolving M2-like macrophages to support vessel remodelling through the paracrine effects of chemokines, metalloproteinases, and growth factors^44,45,64,65^. Aging results in profound alterations in the immune system^66,67^, which may influence arteriogenesis. Here, macrophage mobilization and infiltration was impaired in old compared to young mice. However, these macrophages seemed shifted toward a pro-repair M2 phenotype in old mice, which may highlight the existence of a tissue macrophage-specific compensatory mechanism in old mice. Thus, impaired immune response alone is unlikely to explain the impaired neovascularisation we observe in aging.

Next, focusing on sprouting angiogenesis, we observed a reduction in the proliferative capacity of aged ECs in the ischemic muscle post-HLI, along with diminished aortic sprouting capability. Late passage HUVECs displayed signs of cell cycle arrest and impaired proliferation although we do not observe an upregulation in proteins involved in the canonical induction of cell cycle arrest. This finding is recapitulated in the RNASeq analyses, where we do not observe upregulation of classical markers of cellular senescence p16 and p21 expression. These findings are in opposition to a previous study which described distinct changes in expression of genes regulating cell cycle control and p53-mediated transcription in endothelial cells harvested from 80-year old mesenteric arteries^20^. These differences could be attributed to age differences (our use of 18-months old mice, equivalent to a 56 year old human), or to a difference in EC subtype (our use of ECs harvested from gastrocnemius muscle, which consists mainly of capillary ECs, whereas the described study looked at arterial ECs), or even due to species-related factors that we have not accounted for. In contrast, late passage HUVECs retain their migratory capabilities and their abilities to respond to VEGF, which is in line with our finding that the inhibition of VEGF signalling impacts EC migration, but not proliferation. Overall, our results suggest that the defect in EC proliferation with age is not the result of an impaired VEGF response or the activation of a cell cycle arrest programme.

While undertaking an angiogenic drive, ECs typically upregulate glycolysis for energy and biomass generation^68,69^. While old ECs display an upregulation of pro-angiogenic gene expression, their glycolytic capacity is decreased. We also observed a reduction in ^13^C_6_ enrichment of glycolytic intermediates in late passage HUVECs, indicative of a reduced glycolytic flux. Glycolysis is vital for EC biology, providing over 85% of ATP even in oxygen-replete conditions^70^. We observed a reduction of ^13^C_6_ incorporation into ATP, signifying a reduced level of glucose-derived energy generation, which has important implications in the determination of EC phenotypes (active/quiescent)^70,71^. Glucose metabolism have been shown to directly regulate the cell cycle through lactate^72^, and lactate production was reduced in late passage HUVECs, which may contribute to the defect in proliferation. Glucose catabolism also supports biomass synthesis, diverting intermediates to anabolic pathways like the pentose phosphate pathway (PPP) and serine biosynthesis to support EC proliferation^73,74^. In this study we showed that late passage HUVECs demonstrate elevated gene expression in the PPP, along with normal metabolic flux in PPP and serine biosynthesis, despite compromised glycolytic flux. Thus, old EC may not suffer from impaired biomass synthesis, although further studies are required to properly assess protein homeostasis. Furthermore, we observe a reduction in ^13^C_6_ incorporation into early TCA cycle intermediates (acetyl-CoA and aconitate), which may be due to a reduction in glycolytic flux. Later TCA intermediates (succinic, fumaric, and malic acids) contain a higher proportion of ^13^C_6_ enrichment, which may be the result of disrupted cofactor or enzyme function. From the results of this study, we posit that the glycolysis defect in old ECs may act as a bottleneck for proliferation and neovascularization, due to its centrality in EC function.

In conclusion, we discovered that, while old ECs express elevated levels of pro-angiogenic genes at baseline, they may be ultimately restrained by dysfunctional metabolism, leading to reduced energy production that could prove crucial for cell proliferation. Our future research will focus on identifying ways to boost glycolytic flux in aged ECs and establishing whether glycolysis augmentation could restore proliferation in aged ECs. Clarifying these mechanisms will help us understand the age-associated decline in EC, with crucial implications for the maintenance of tissue perfusion and capability to respond to ischaemic diseases.

## FIGURES AND LEGENDS

**Fig. S1:**
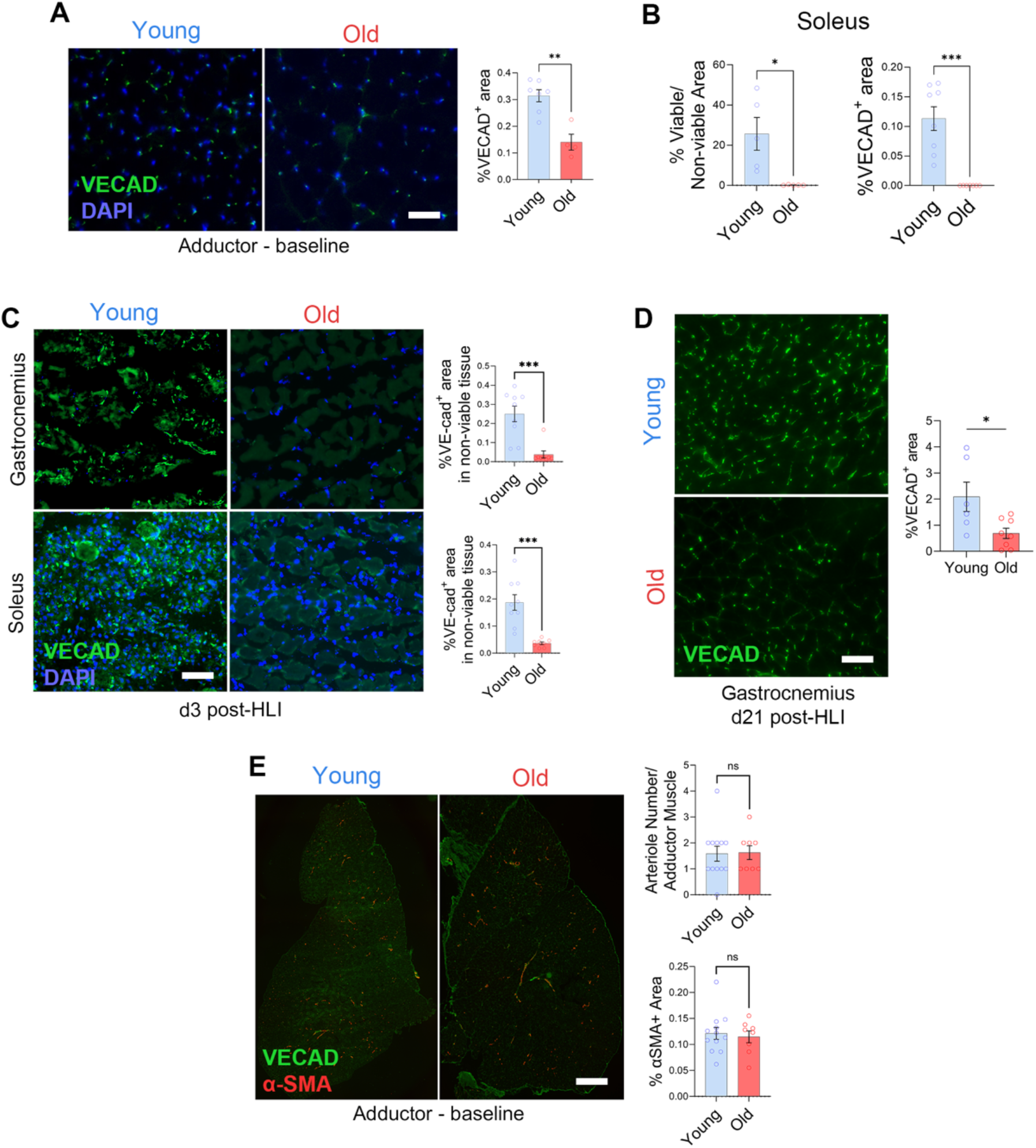
Aged mice undergo slower neovascularisation and suffer more muscle damage post-hindlimb ischemia. **(A)** Adductor muscle microvasculature at baseline as depicted by representative images (left) and quantification (right) of transverse sections stained with VE-cad. n=8-12 per group. Scale bar represents 50µm. **(B)** Quantification of ischemic soleus muscle microvasculature at 3 days post-HLI of transverse sections stained with VE-cad. n=8-9 per group. Scale bar represents 50µm. **(C)** Microvasculature within the ischemic regions of gastrocnemius and soleus muscle at 3 days post-HLI as depicted by representative images (left) and quantification (right) of transverse sections stained with VE-cad. n=8-9 per group. Scale bar represents 50µm. **(D)** Quantification of gastrocnemius muscle microvasculature at 21 days post-HLI of transverse sections stained with VE-cad. n=8-9 per group. Scale bar represents 50µm. **(E)** Arterioles in baseline adductor muscle as depicted by representative images (left) and quantification (right) of transverse sections stained with VE-cad and α-SMA. n=8-12 per group. Scale bar represents 50µm. Data are expressed as mean ± S.E.M. ** p≤0.01, *** p≤0.001 and **** p≤0.0001 by Student’s t-test.

**Fig. S2:**
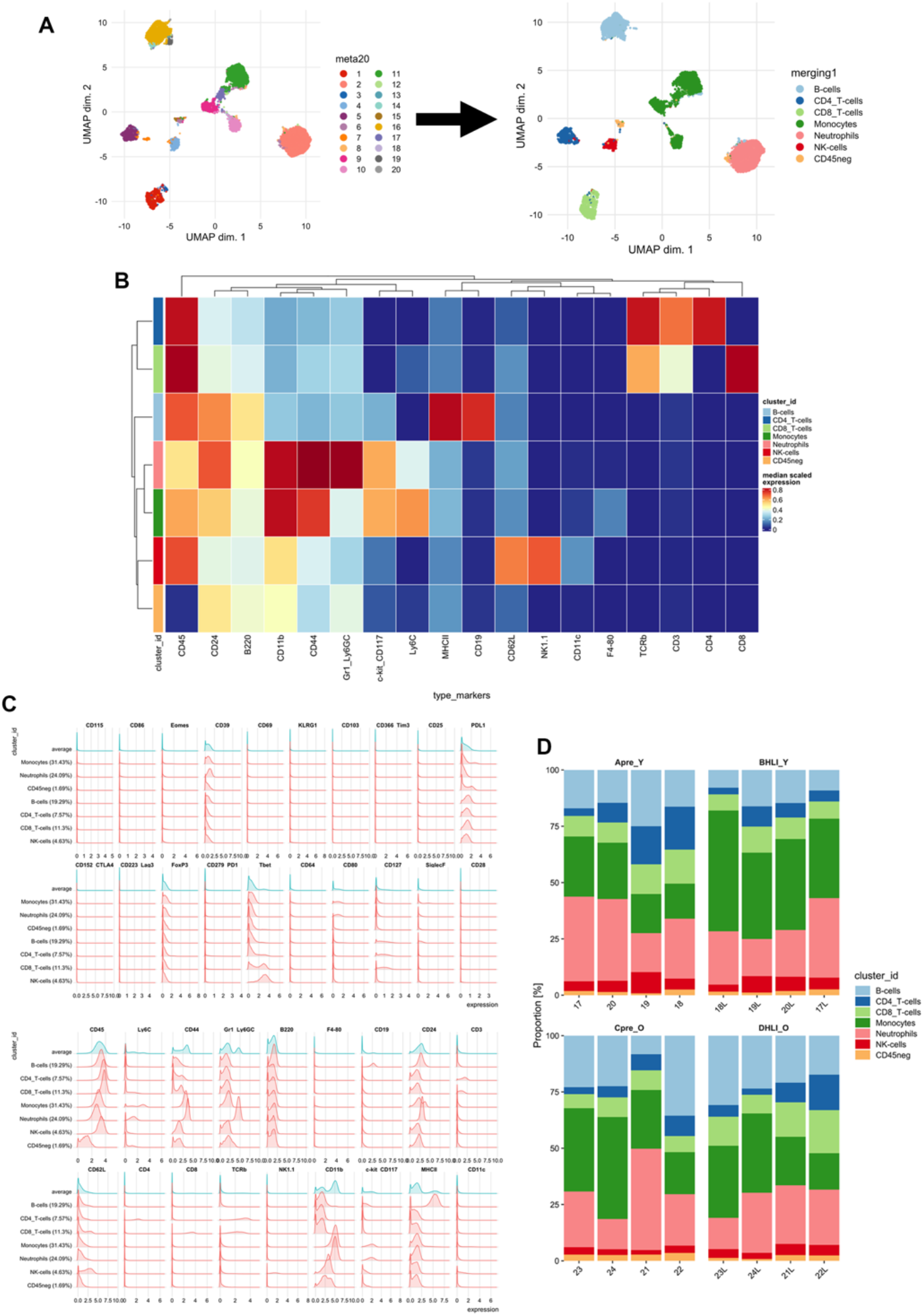
CyTOF analysis on immune recruitment post-hindlimb ischemia in young and old mice. **(A)** UMAP plot for the PBMC dataset, where cells are coloured according to the manual merging of the 20 cell populations, obtained with FlowSOM, into 7 PBMC populations. **(B)** Heatmap of the median marker intensities of the 7 PBMC populations obtained with FlowSOM. **(C)** Distributions of marker intensities (arcsinh-transformed) of the 7 PBMC populations obtained with FlowSOM. **(D)** Relative abundance of the 7 PBMC populations in each sample (x-axis), segregated by age group (young and old) and condition (pre- and post-HLI), represented by barplots.

**Fig. S3:**
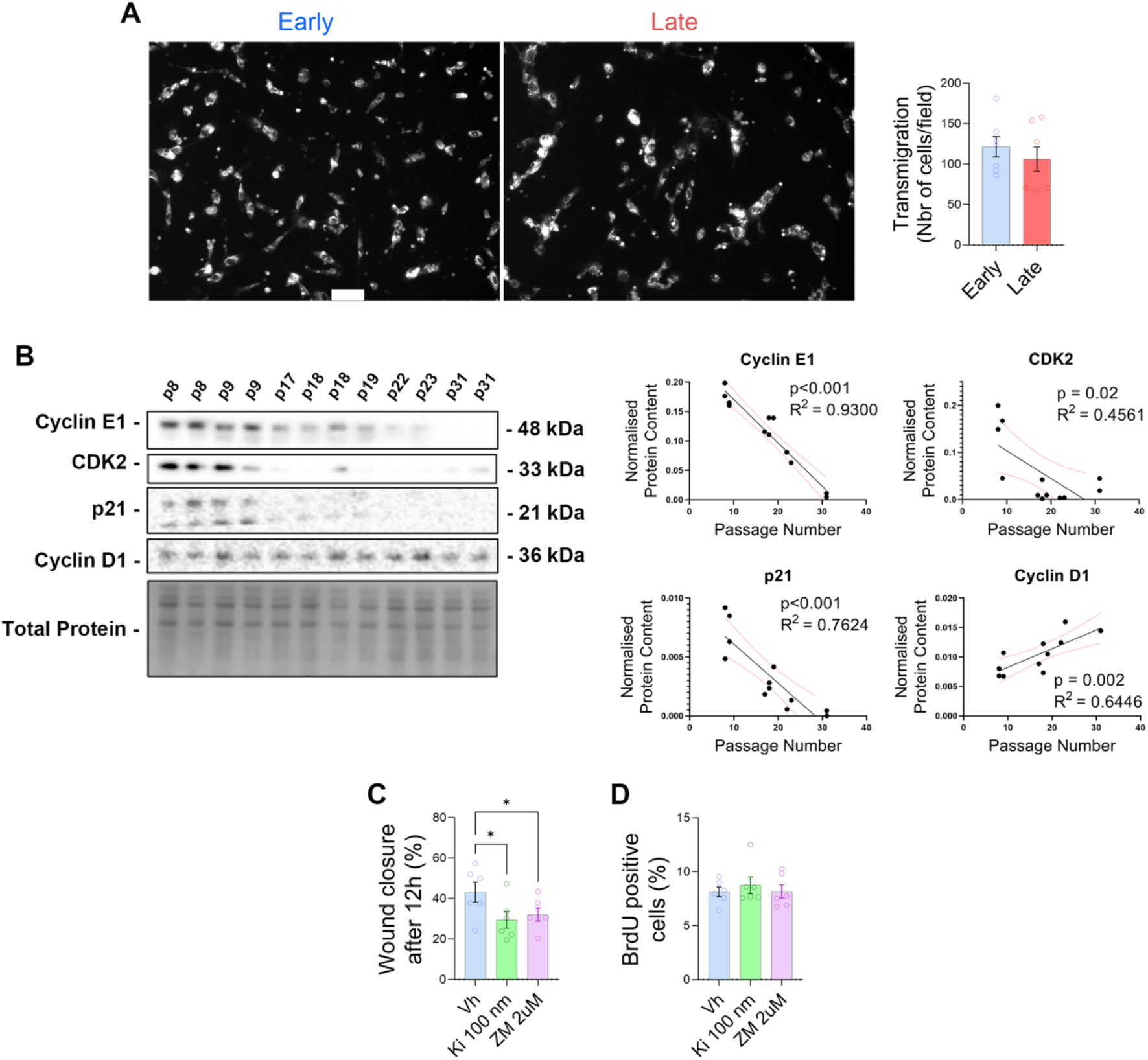
Impairment of EC proliferation results in neovascularisation defect in old mice. **(A)** Early- and late-passage HUVEC transmigration through the polycarbonate membrane of a Boyden chamber from unsupplemented EBM-2 medium towards supplemented EGM-2 medium. Scale bar represents 50 μm. **(B)** Western blot of cell-cycle proteins on lysates of early and late passage HUVECs. All proteins normalized to total protein stain. **(C-D)** Quantification of (C) EC migration across a scratch and (D) BrdU incorporation assay of early passage HUVECs treated with the VEGF inhibitors KI8751 (100 nM) and ZM323881 (2 µM). n=6 per group.

## Notes

### Competing Interest Statement

The authors have declared no competing interest.

