## Supplementary Tables for "Impaired glycolysis in aged endothelial cells is associated with reduced neovascularisation upon tissue ischemia"

**Supplementary Table S1: Antibodies**

| Target antigen | Vendor | Catalog # | Working concentration |
|---|---|---|---|
| Laminin | Sigma | L9393 | 1/200 (IHC) |
| VE-Cadherin | BD Pharmingen | 555289 | 1/200 (IHC) |
| CD45 | BD Biosciences | 550539 | 1:200 |
| CD68 | Biorad | MCA1957T | 1/500 (IHC) |
| HO-1 | Abcam | 13243 | 1/200 (IHC) |
| ERG | Cell Signaling Technology | #4695 | 1/100 (IHC) |
| BrdU | BD Biosciences | 555627 | 1/200 (ICC) |
| α-SMA | Cell Signaling Technology | #19245 | 1/500 (IHC) |
| Goat anti-Rabbit Alexa Fluor 680 | Thermo Fisher Scientific | A21109 | 1/500 (IHC) |
| Goat anti-Rabbit Alexa Fluor 405 | Thermo Fisher Scientific | A31556 | 1/500 (IHC) |
| Goat anti-Rat Alexa Fluor 488 | Thermo Fisher Scientific | A11006 | 1/500 (IHC) |
| Donkey anti-Rabbit Alexa Fluor 488 | Thermo Fisher Scientific | A21206 | 1/500 (IHC) |
| Cyclin E1 | Cell Signaling Technology | 20808 | 1/1000 (WB) |
| CDK2 | Cell Signaling Technology | 2546 | 1/1000 (WB) |
| p21 | Cell Signaling Technology | 2947 | 1/1000 (WB) |
| Cyclin D1 | Cell Signaling Technology | 2978 | 1/1000 (WB) |
| pVEGFR2 | Cell Signaling Technology | 2478S | 1/1000 (WB) |
| VEGFR2 | Cell Signaling Technology | 2479 | 1/1000 (WB) |
| pERK | Cell Signaling Technology | 4370 | 1/2000 (WB) |
| ERK | Cell Signaling Technology | 4695 | 1/1000 (WB) |
| p-p38 | Cell Signaling Technology | 9211 | 1/1000 (WB) |
| p38 | Cell Signaling Technology | 9212 | 1/1000 (WB) |
| Anti-Rabbit HRPO | Thermo Fisher Scientific | 31460 | 1/20000 (WB) |
| Anti-mouse HRPO | Jackson ImmunoResearch Labs | 115-035-146 | 1/15000 (WB) |

**Commented [k1]:** Should I add other cell cycle Western Abs that we didn't include in the figure?

**Commented [FA2R1]:** You only put in the antibodies shown in the data

**Supplemental table S2: DNA oligo primers**

| Primer Name | Gene | Sequence |
|---|---|---|
| Arg1 | Arginase | Fw: AACCAGCTCTGGGAATCTGC<br>Rv: TCCTGGTACATCTGGGAACTTT |
| Ym1 | Chitinase-like 3 | Fw: AGAAGGGAGTTTCAAACCTGGT<br>Rv: GTCTTGCTCATGTGTGTAAGTGA |
| IL10 | Interleukin 10 | Fw: GCTGTCATCGATTTCTCCCCT<br>Rv: GACACCTTGGTCTTGGAGCTTAT |
| Mrc1 | Mannose Receptor C-Type 1 | Fw: GAGGCTGATTACGAGCAGTG<br>Rv: TTGGTTCACCGTAAGCCCAAT |
| Tnfα | Tumor necrosis factor alpha | Fw: ACGGCATGGATCTCAAAGAC<br>Rv: AGATAGCAAATCGGCTGACG |
| Ccl2 | Chemokine (C-C motif) ligand 2 | Fw: CAGGTCCCTGTCATGCTTCT<br>Rv: GCGTTAACTGCATCTGGCTGA |
| Cxcl1 | Chemokine (C-X-C motif) ligand 1 | Fw: CCAGAGCTTGAAGGTGTTGC<br>Rv: CCATTCTTGAGTGTGGCTATGAC |
| IL1b | Interleukin 1 beta | Fw: GTGTCTGAAGCAGCTATGGCA<br>Rv: CAGGTCATTCTCATCACTGTCAA |
